# A p53-dependent FBXO44-RAD18 axis limits mutagenesis by terminating translesion DNA synthesis

**DOI:** 10.64898/2026.07.27.739831

**Authors:** Alessio Butera, Sabrina Caporali, Francesco Capradossi, Paul Smith, Nino Tabagari, Nora Nikoloska, Olga Mayans, Ana Janic, Andreas J. Gruber, Vincenzo D’Angiolella, Ivano Amelio

## Abstract

DNA lesions continually challenge genome replication and threaten genome integrity. DNA damage tolerance pathways, including translesion DNA synthesis (TLS), allow cells to bypass lesions and prevent stalled forks from collapsing into double-strand breaks. Because TLS polymerases are intrinsically error-prone, however, this pathway must be tightly restrained; persistent or deregulated TLS can increase mutagenesis, create therapeutic vulnerabilities, and promote aggressive cancer phenotypes.

Through integrated transcriptional profiling, genome-wide CRISPR/Cas9 screening for replication-stress sensitivity, and complementary proteomic analyses, we identify F-box protein 44 (FBXO44) as a late p53-responsive regulator of the TLS mediator RAD18. FBXO44 promotes RAD18 ubiquitination during recovery from replication stress and facilitates shutdown of RAD18-dependent PCNA monoubiquitination. Consistently, FBXO44 loss delays resolution of replication stress and TLS signaling, increases mutation frequency, and is associated with elevated mutational burden and therapy resistance in experimental models and patient datasets.

These findings define a p53-FBXO44-RAD18 regulatory axis that limits mutagenic TLS and helps safeguard genome integrity after replication stress.

## Introduction

Faithful genome duplication is essential for accurate genetic inheritance^1,2^. Replication is continually challenged by endogenous and exogenous stressors that compromise fork progression and can generate double-strand breaks, key drivers of genomic instability^3^. To preserve genome integrity, eukaryotic cells deploy DNA damage tolerance (DDT) pathways that allow replication to continue across damaged templates and help ensure completion of genome duplication^2,4,5^.

Translesion DNA synthesis (TLS) is a direct lesion-bypass mechanism in which specialized polymerases replicate across damaged DNA^6^. Several chemotherapeutic agents, including platinum-based compounds, activate TLS. However, because TLS polymerases have intrinsically low fidelity, insufficient control of TLS can be exploited by cancer cells to increase mutagenesis, accelerate tumor evolution, and acquire therapy resistance^7–10^.

TLS initiation is controlled in part by the replication-stress kinase ataxia telangiectasia and Rad3-related (ATR), which promotes activation of the E3 ubiquitin ligase RAD18. RAD18 is recruited to stalled forks, where it monoubiquitinates PCNA on lysine 164 and thereby enables recruitment of TLS polymerases for lesion bypass. Once bypass is complete, TLS must be terminated promptly to limit mutational risk. Although RAD18 activation has been extensively investigated, how RAD18 is restrained during TLS termination remains poorly understood^11–14^.

The tumor suppressor p53 is a central regulator of the cellular response to replication stress and DTT, including TLS. Loss of p53 function can increase replication stress, alter TLS and dependence on ATR-mediated signaling, thereby creating vulnerabilities to ATR inhibition. We therefore reasoned that p53-regulted genes might determine how cells regulate TLS and tolerate impaired ATR signaling.

To identify p53-linked regulators of the replication-stress response, we integrated p53-associated transcriptional profiles and predicted chromatin-binding sites with a genome-wide CRISPR/Cas9 screen for sensitivity to ATR inhibition. This strategy nominated the F-box protein FBXO44. Proximity proteomics and differential ubiquitination profiling then connected FBXO44 to RAD18 regulation. We found that FBXO44 deficiency reduces RAD18 ubiquitination, delays resolution of replication stress, and sustains PCNA monoubiquitination at lysine 164. Persistent TLS signaling is accompanied by increased mutation frequency and resistance to genotoxic therapy, while clinical analyses associate FBXO44 expression with TP53 status, tumor mutational burden, therapy response, and disease-free survival.

Together, these data reveal a p53-linked mechanism that restrains RAD18 during TLS termination and limits mutation-associated genome instability.

## Results

### Integrative analysis identifies FBXO44 as a determinant of ATR-inhibitor sensitivity

ATR-dependent signaling helps cells manage replication-fork challenges and complete DNA replication. Because TP53 mutations create replication-stress vulnerabilities^15,16^, we designed an integrative strategy to identify p53-associated genes that influence the response to ATR inhibition. We combined p53-dependent transcriptional profiles^15,17,18^ and predicted chromatin-binding sites^19^ with a genome-wide CRISPR/Cas9 screen^20^ using the ATR inhibitor AZD6738 (Fig. 1A). Alongside SETD2 and FANCC, two established regulators of the replication stress response^21–24^, FBXO44 emerged as a top candidate, and its depletion was associated with synthetic lethality under ATR inhibition (Fig. 1B). Independent validation in retinal pigment epithelium (RPE) cells confirmed increased sensitivity to AZD6738 after endogenous FBXO44 depletion. Three independent siRNAs targeting FBXO44 consistently reduced clonogenic survival after AZD6738 treatment (Fig. 1C and Suppl. Fig. 1A).

**Figure 1.**
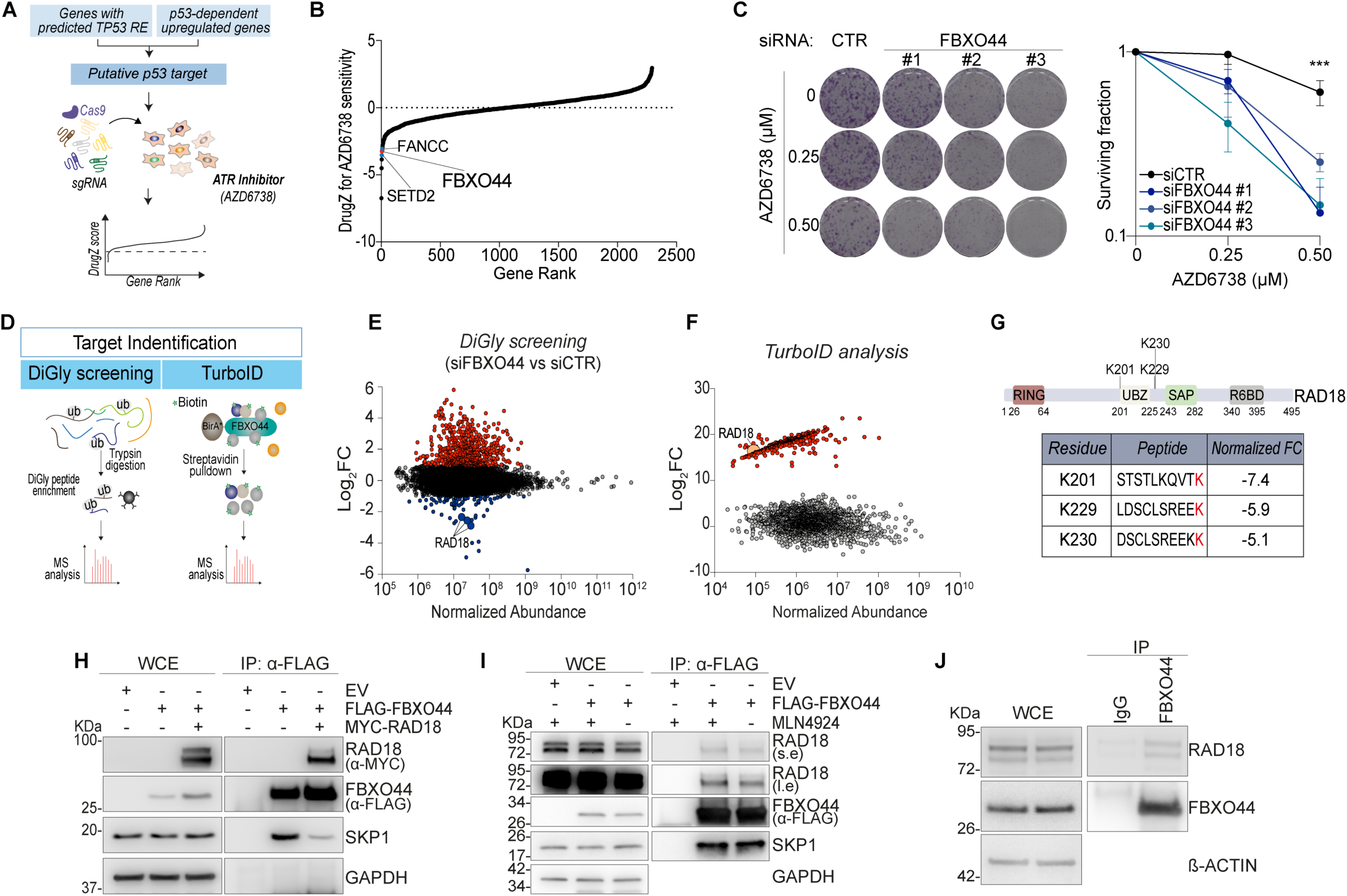
FBXO44 is a novel regulator of the replication stress response. A) Integrative multistep strategy used to identify genetic determinants of the replication-stress response. B) DrugZ sensitivity plot identifying FBXO44 among the top five hits associated with synthetic lethality after ATR inhibition (AZD6738) in RPE p53^-/-^. C) Clonogenic assay in RPE p53^-/-^ cells after FBXO44 depletion and AZD6738 treatment. N = 3 independent biological replicates; ***p < 0.001. Bars indicate standard deviation. P values were calculated by ordinary one-way ANOVA with Tukey’s correction. D) Strategy used to identify candidate FBXO44 targets. E, F) Log2FC plots for diGly profiling (E) and TurboID (F). G) RAD18 domain schematic indicating peptides and lysine residues with reduced ubiquitination after FBXO44 depletion. H-J) Co-immunoprecipitation assays showing interaction between RAD18 and FBXO44. N = 3 independent biological replicates; one representative experiment is shown.

These findings identify FBXO44 as a candidate modifier of ATR-inhibitor sensitivity.

### RAD18 is a putative FBXO44 substrate

Because FBXO44 is poorly characterized and its substrates remain largely unknown, we used two complementary mass-spectrometry-based approaches. We performed di-glycine (diGly) profiling to enrich peptides carrying ubiquitin-remnant modifications and TurboID proximity labeling to define the FBXO44-proximal proteome (Fig. 1D). Integrating these datasets identified the ubiquitin E3 ligase RAD18 as a candidate FBXO44 substrate or proximal interactor (Fig. 1E, F).

DiGly profiling revealed three RAD18 lysines with reduced ubiquitination after FBXO44 depletion: K201 (fold change, -7.4), K229 (fold change, -5.9), and K230 (fold change, -5.1) (Fig. 1G). K201 has previously been linked to RAD18 self-inhibition^25^, whereas K229 and K230 are, to our knowledge, uncharacterized ubiquitination sites (Suppl. Fig. 1B, C). We then validated the FBXO44-RAD18 interaction by co-immunoprecipitation under multiple conditions, including endogenous and exogenous expression systems (Fig. 1H-J and Suppl. Fig. 1D). Together, these findings support RAD18 as a putative FBXO44 substrate.

### FBXO44 promotes RAD18 ubiquitination

RAD18 is activated in response to perturbed DNA replication^25–28^. We therefore asked whether replication stress modulates the FBXO44-RAD18 interaction. After inducing replication stress with the ribonucleotide reductase inhibitor hydroxyurea (HU)^29^, we observed enhanced FBXO44-RAD18 binding, with the strongest interaction during recovery after HU withdrawal (Fig. 2A). Thus, replication-stress signaling promotes FBXO44 association with RAD18.

**Figure 2.**
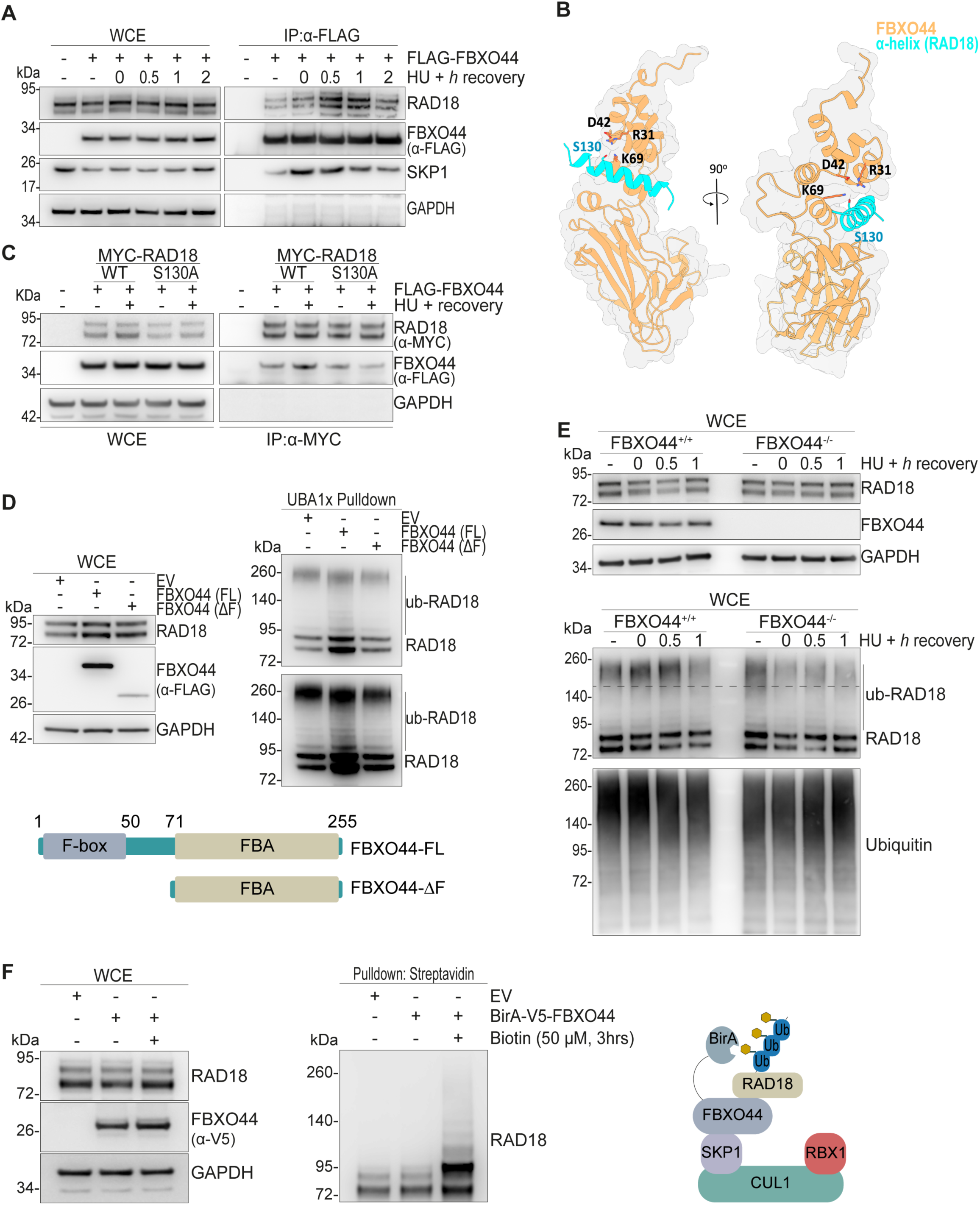
FBXO44 promotes RAD18 ubiquitination. A) Co-immunoprecipitation showing that replication stress enhances FBXO44-RAD18 interaction in HEK293T cells; one representative experiment of three independent biological replicates is shown. B) AlphaFold3 model of the complex formed by FBXO44 (gold) and a RAD18 helical segment (cyan; residues 123-144). RAD18 S130 is buried in the interface facing a charged patch. C) Co-immunoprecipitation showing that RAD18 S130A loses the replication-stress-associated increase in FBXO44 binding. D) UBE pull-down ubiquitination assay showing that full-length FBXO44, but not the DeltaF mutant, promotes RAD18 ubiquitination; one representative experiment of three independent biological replicates is shown. E) UBE pull-down under replication stress showing increased high-molecular-weight ubiquitinated RAD18 in FBXO44-proficient (FBXO44+/+) cells but not in FBXO44- deficient (FBXO44-/-) cells. Ubiquitin was probed for pull-down normalization; one representative experiment of three independent biological replicates is shown. F) E-STUB assay showing increased biotin labeling of ubiquitinated RAD18; one representative experiment of three independent biological replicates is shown.

To explore the structural basis of FBXO44-RAD18 binding, we modeled the complex in silico using AlphaFold3^30^ and HDOCK^31^. The models predicted that RAD18 engages FBXO44 through a 21-amino-acid segment, 122-RQSLKQGSRLMDNFLIREMSG-143, that forms an approximately amphipathic alpha helix within a long intrinsically disordered region (Fig. 2B and Suppl. Fig. 2). On FBXO44, the interface involves the hinge between the N-terminal helical domain and the C-terminal beta sandwich: helices 2, 3, and 5 of the N-terminal domain and helix 7 of the beta sandwich contribute to RAD18 binding. Within this interface, RAD18 S130 is buried and oriented toward a charged patch formed by K69, R31, and D42 of FBXO44.

To test this model, we generated a RAD18 S130A mutant and performed co-immunoprecipitation assays. Consistent with the prediction, RAD18 S130A failed to show the replication-stress-recovery-associated increase in FBXO44 binding observed with wild-type RAD18 (Fig. 2C).

F-box proteins typically recruit substrates to SCF ubiquitin ligase complexes^32^. To assess whether FBXO44 promotes RAD18 ubiquitination, we performed in vivo ubiquitination assays using ubiquitin binding entities (UBE) pull-down. Unbiased enrichment of ubiquitinated proteins showed that full-length FBXO44 (FL), but not an F-box-deleted mutant (DeltaF) unable to assemble into an SCF complex, induced high-molecular-weight RAD18 species consistent with ubiquitination (Fig. 2D).

Because FBXO44-RAD18 binding peaked during recovery from replication stress, we next examined RAD18 ubiquitination kinetics after HU treatment and withdrawal. RAD18 ubiquitination increased after HU treatment and peaked during recovery in FBXO44- proficient cells, whereas this induction was abolished in FBXO44-deficient cells (Fig. 2E). To further support FBXO44-mediated RAD18 ubiquitination, we used E3 substrate tagging by ubiquitin biotinylation (E-STUB)^33^, a proximity-based assay for identifying cellular E3 ligase substrates. After ectopic expression of BirA-V5-FBXO44 and ubiquitin containing a biotin acceptor peptide, ubiquitinated RAD18 was detected after streptavidin pull-down (Fig. 2F). These data support FBXO44-dependent RAD18 ubiquitination in cells.

Together, these results identify FBXO44 as a cellular regulator of RAD18 ubiquitination.

### FBXO44 loss compromises TLS termination

To assess the functional consequences of FBXO44-mediated RAD18 ubiquitination, we analyzed RAD18 protein abundance and half-life under basal and replication-stress conditions. FBXO44 did not measurably promote RAD18 degradation (Suppl. Fig. 3A-D). Although SCF E3 ligases are classically associated with proteasomal degradation^32^, F-box proteins can also mediate non-proteolytic ubiquitination events that regulate substrate activity, localization, or interactions^34–36^. We therefore examined how FBXO44 affects RAD18 activity and function.

During replication of damaged templates, RAD18 monoubiquitinates PCNA at lysine 164 (PCNA-mUb), an established marker of TLS activation^37,38^. We therefore monitored PCNA-mUb as a readout of RAD18 catalytic activity during induction and resolution of replication stress after HU treatment or UV-C irradiation. FBXO44-deficient cells showed delayed resolution of PCNA-mUb during recovery, consistent with defective TLS termination (Fig. 3A-D).

**Figure 3.**
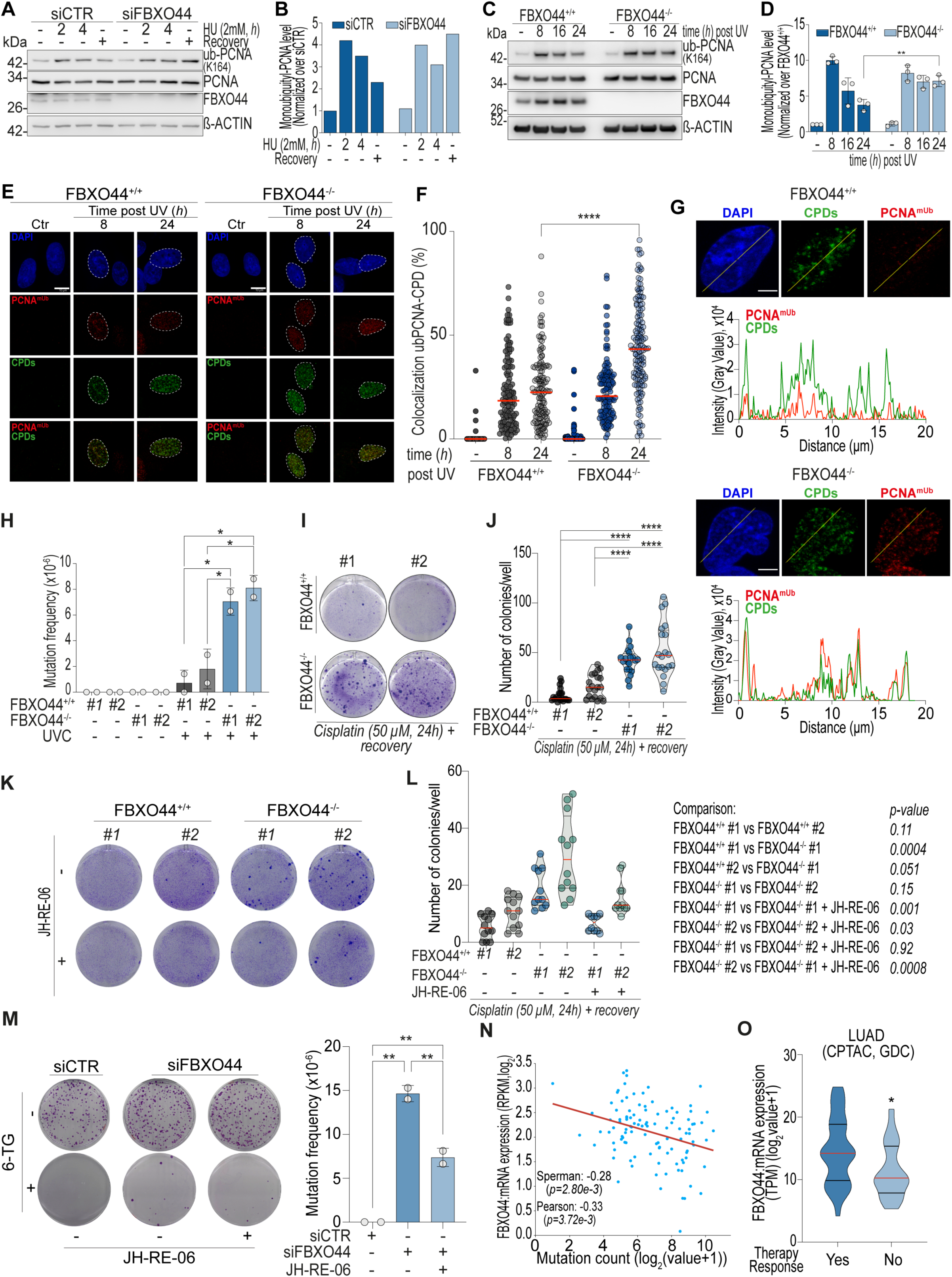
FBXO44 loss delays resolution of replication stress and PCNA monoubiquitination and is associated with mutagenesis and chemoresistance. A, B) Immunoblot (A) and quantification (B) showing delayed resolution of monoubiquitinated PCNA (K164) in FBXO44-silenced A549 cells after hydroxyurea (HU)-induced replication stress; one representative experiment of three independent biological replicates is shown. C, D) Immunoblot (C) and quantification (D) showing delayed resolution of monoubiquitinated PCNA (K164) after FBXO44 knockout and UV-C-induced replication stress. Histograms show mean +/- standard deviation from N = 3 independent replicates; **p < 0.01. P values were calculated by ordinary one-way ANOVA with Tukey’s correction. E, F) Immunofluorescence (E) and quantification (F) showing PCNA-mUb colocalization with UV-C-induced CPD lesions. In F, red bars indicate the mean. Scale bar, 10 µm; ****p < 0.0001. P values were calculated by ordinary one-way ANOVA with Tukey’s correction. G) Subnuclear analysis of PCNA-mUb and CPDs. Left: representative images; scale bar, 5 µm. Right: fluorescence intensity histograms along the yellow line in the corresponding image. H) In vitro HPRT mutagenesis assay showing increased mutation frequency in A549 FBXO44-/- clones after UV-C damage. Histograms show mean +/- standard deviation from two independent experiments. Two wild- type (FBXO44+/+) and two FBXO44-knockout (FBXO44-/-) clones were used; *p < 0.05. P values were calculated by ordinary one-way ANOVA with Tukey’s correction. I, J) Colony survival assay after cisplatin treatment (50 µM, 24 h) showing increased resistance of FBXO44-/- clones. Violin plots (J) show median (red line) and quartiles (black dashed line) from N = 5 independent replicates; ****p < 0.0001. P values were calculated by ordinary one-way ANOVA with Tukey’s correction. K, L) Micrograph (K) and quantification (L) of colony survival after cisplatin treatment with or without the REV1 inhibitor JH-RE-06. N = 4 independent biological replicates. P values were calculated by ordinary one-way ANOVA with Dunnett’s correction. M) HPRT mutagenesis assay in U2OS cells proficient or deficient for FBXO44 after cisplatin (5 µM, 24 h) and JH-RE-06 (2 µM, 24 h) treatment. Histograms show mean +/- standard deviation from two independent experiments; **p < 0.01. P values were calculated by ordinary one-way ANOVA with Sidak’s correction. N) Correlation between FBXO44 mRNA expression and mutation count in lung adenocarcinoma (LUAD). Spearman: -0.28, p = 2.80e-3; Pearson: -0.33, p = 3.72e-3. O) FBXO44 mRNA expression in lung adenocarcinomas stratified by therapy response; *p < 0.05. P values were calculated by unpaired t-test.

To validate these findings at damage-associated foci, we used confocal microscopy to quantify PCNA-mUb at cyclobutane pyrimidine dimer (CPD) lesions after UV-C irradiation. Consistent with the immunoblot data, PCNA-mUb colocalization with CPD lesions persisted at later time points in FBXO44-/- cells compared with FBXO44+/+ controls (Fig. 3E, F). Subnuclear analysis 24 h after UV-C irradiation further showed reduced PCNA-mUb at CPD lesions in FBXO44+/+ cells, whereas this decrease was not observed in FBXO44-/- cells (Fig. 3G).

These data indicate that FBXO44 loss delays shutdown of RAD18-dependent PCNA monoubiquitination during TLS termination.

### FBXO44 loss promotes mutagenesis and chemoresistance

TLS polymerases are error-prone because they lack proofreading activity and have flexible catalytic sites that accommodate damaged DNA^39^. Because FBXO44 loss sustained PCNA- mUb after replication stress, we asked whether FBXO44 deficiency increases mutation frequency. We used the HPRT assay, an established in vitro method that measures mutagenic inactivation of the X-linked HPRT gene by selecting for survival in the presence of a toxic nucleotide analogue^32,40^ in lung adenocarcinoma cells (A549). After UV-C-induced TLS activation, A549 FBXO44-/- clones showed significantly increased mutation frequency compared with isogenic FBXO44+/+ controls (Fig. 3H and Suppl. Fig. 4A).

Mutagenesis can support cancer-cell survival and evolution under therapeutic stress, contributing to relapse after treatment^6,7^. Platinum-based agents such as cisplatin activate TLS, and lung cancer cells can exploit this pathway to acquire mutations that promote cisplatin resistance^8,9^. We therefore tested whether FBXO44 loss gives lung cancer cells a selective advantage during acquisition of cisplatin resistance. A549 cells were exposed to high-dose cisplatin, which produced an initially lethal response (>90% cell death), and residual resistant colonies were monitored for 4 weeks. FBXO44 knockout substantially increased colony formation compared with FBXO44 wild-type clones (Fig. 3I, J and Suppl. Fig. 4B).

To test whether this phenotype depends on TLS, we treated cells with the TLS Polymerase inhibitor JH-RE-06^41^. JH-RE-06 reduced the increased colony formation observed in FBXO44-deficient cells (Fig. 3K, L). In U2OS osteosarcoma cells, endogenous FBXO44 depletion also increased mutational burden after cisplatin treatment, and this effect was again reduced by JH-RE-06 (Fig. 3M and Suppl. Fig. 4C). Consistent with the experimental data, in lung adenocarcinoma patient cohorts, lower FBXO44 expression was associated with higher tumor mutational burden and poorer therapy response (Fig. 3N, O). Low FBXO44 expression was also associated with disease progression or recurrence and shorter disease-free survival (Suppl. Fig. 4D-G).

Overall, FBXO44 loss promotes mutagenesis and cisplatin resistance in cellular models, and low FBXO44 expression is associated with adverse post-treatment clinical features.

### FBXO44 is a late p53-responsive gene

Finally, to connect the FBXO44-RAD18 axis with tumor-suppressive signaling and the initial screening strategy (Fig. 1A), we tested whether FBXO44 is regulated by p53-dependent transcriptional programs. Activation of p53 with the MDM2 inhibitor Nutlin-3a induced FBXO44 mRNA and protein after 6 h in p53-proficient cells, but not in p53-deficient or p53- mutant cell lines (Fig. 4A and Suppl. Fig. 5A-D). Similar induction was observed after treatment with replication-stress- or TLS-activating genotoxic stimuli, including UV-C irradiation and HU (Fig. 4B, C and Suppl. Fig. 6A-D). These results identify FBXO44 as a p53-dependent DNA-damage-responsive gene.

**Figure 4.**
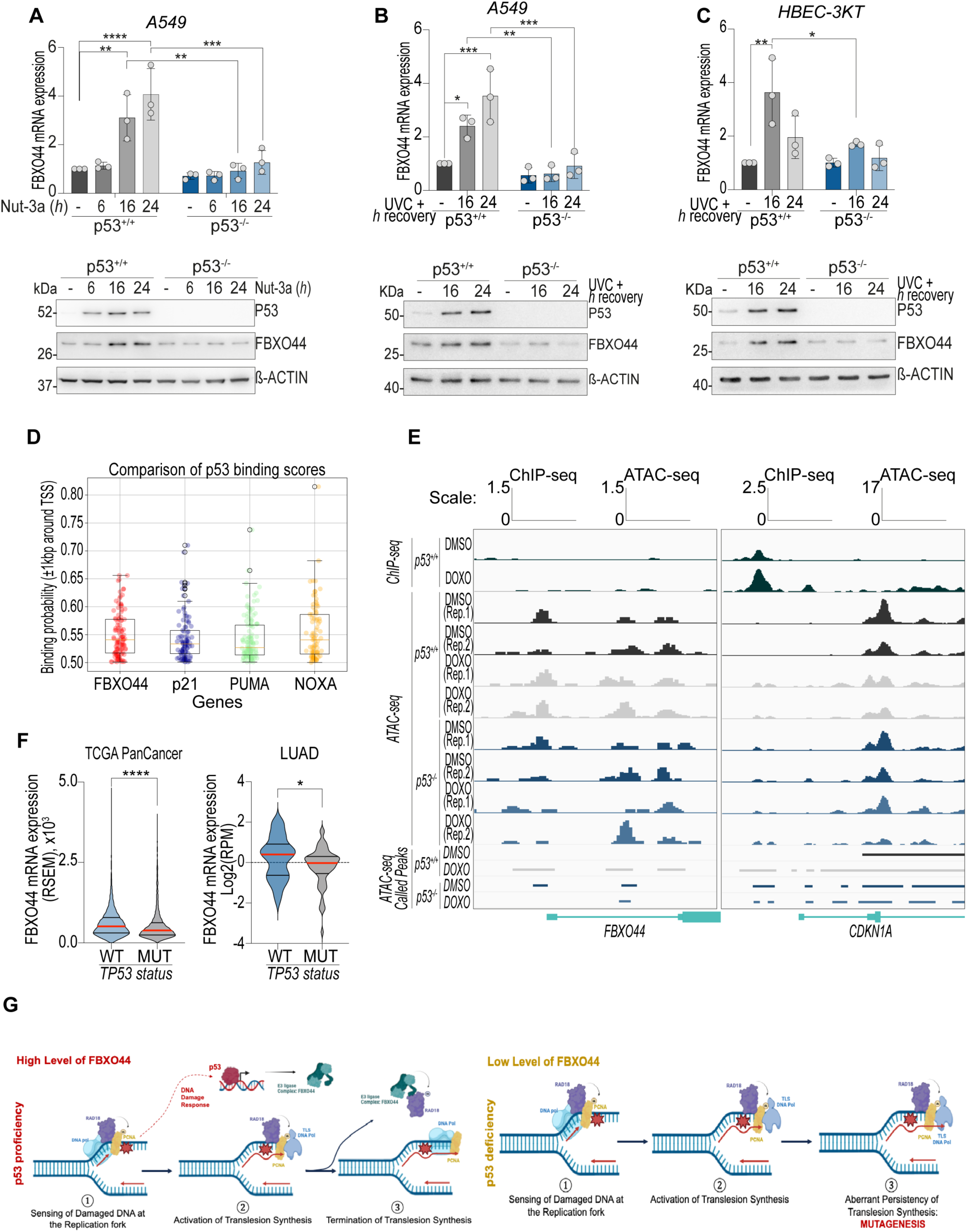
FBXO44 is a late p53-responsive target gene. A) RT-qPCR (upper) and immunoblot (lower) showing FBXO44 upregulation kinetics after p53 activation with the MDM2 inhibitor Nutlin-3a in A549 cells proficient (p53+/+) or deficient (p53-/-) for p53. Bars represent mean +/- standard deviation from three independent biological replicates; **p < 0.01, ***p < 0.001, ****p < 0.0001. P values were calculated by ordinary one-way ANOVA with Sidak’s correction. B) RT-qPCR (upper) and immunoblot (lower) showing FBXO44 levels after UV-C irradiation in A549 p53+/+ and p53-/- cells. Bars represent mean +/- standard deviation from three independent biological replicates; *p < 0.05, **p < 0.01, ***p < 0.001. P values were calculated by ordinary one-way ANOVA with Sidak’s correction. C) RT-qPCR (upper) and immunoblot (lower) showing FBXO44 levels after UV-C irradiation in HBEC-3KT p53+/+ and p53-/- cells. Bars represent mean +/- standard deviation from three independent biological replicates; *p < 0.05, **p < 0.01. P values were calculated by ordinary one-way ANOVA with Sidak’s correction. D) p53-binding probability analysis across the FBXO44 promoter region. P21, PUMA, and NOXA were used as positive controls. E) ATAC-seq chromatin-accessibility peaks and ChIP-seq tracks showing increased accessibility at the FBXO44 promoter in p53-proficient (p53+/+) cells without detectable p53 binding. CDKN1A is shown as a direct p53 target. F) FBXO44 expression according to TP53 status across cancer types (PanCancer) and in LUAD. Data source: cBioPortal. G) Working model for the p53- FBXO44-RAD18 axis in genome-integrity maintenance. In p53-proficient cells, replication stress activates TLS and p53 signaling. p53-dependent FBXO44 upregulation contributes to timely TLS termination by restraining RAD18. Upon p53 loss, FBXO44-mediated control of RAD18 is reduced, leading to persistent TLS, mutagenesis, and chemoresistance.

FBXO44 induction occurred only after sustained p53 activation (>6 h), suggesting that p53 may regulate FBXO44 indirectly. Consistent with this model, comparative analysis of predicted p53 binding sites across the FBXO44 locus and established p53 target genes indicated a lower likelihood of direct p53 binding at FBXO44 (Fig. 4D). ATAC-seq analysis showed increased chromatin accessibility at the FBXO44 promoter in p53+/+ but not p53-/- cells, whereas ChIP-seq did not detect p53 occupancy at this locus (Fig. 4E). Thus, FBXO44 appears to be a late p53-responsive gene, likely regulated through secondary chromatin remodeling events associated with sustained p53 activation rather than direct p53 binding^17^.

We next examined the clinical relevance of the p53-FBXO44 axis in patient datasets. Consistent with the experimental data, FBXO44 expression correlated with TP53 status and was higher in TP53 wild-type tumors than in TP53-mutant cohorts (Fig. 4F). Overall, these data place FBXO44 within a p53-associated tumor-suppressive gene network.

## Discussion

Our study identifies FBXO44 as a p53-responsive regulator of mutagenic TLS termination. After replication stress, p53-dependent signaling induces FBXO44, which constrains RAD18, a central TLS regulator. RAD18 monoubiquitinates PCNA to recruit TLS polymerases and initiate error-prone lesion bypass; our data suggest that FBXO44 helps turn this signal off during recovery, thereby limiting mutagenesis.

TLS termination has been linked primarily to USP1, the PCNA-directed deubiquitinase that removes ubiquitin from PCNA lysine 164^42–44^. That step reduces TLS polymerase recruitment, but it does not fully explain how RAD18 itself is restrained once lesion bypass is complete. Here, we show that FBXO44 binds RAD18 and promotes RAD18 ubiquitination during recovery from replication stress. FBXO44 loss sustains PCNA monoubiquitination after HU or UV-C treatment, indicating persistent RAD18-dependent TLS signaling. A model emerging from these findings is that FBXO44-mediated, non-proteolytic ubiquitination limits RAD18 activity, enabling PCNA deubiquitination and TLS shutdown. When this control is lost, low-fidelity polymerases may remain engaged for longer, increasing mutation frequency^39^. Consistent with this model, FBXO44-deficient cells accumulate more mutations after UV-C irradiation and display increased cisplatin resistance (Fig. 4G).

The delayed kinetics of FBXO44 induction further support a role in resolution rather than initiation of the replication-stress response. Promoter scanning, ATAC-seq, and ChIP-seq analyses did not provide evidence for direct p53 binding at the FBXO44 locus. Instead, the data are consistent with an indirect p53-dependent transcriptional response, potentially downstream of p53-driven chromatin remodeling^17^. Such timing may allow cells to activate TLS rapidly when forks stall, while inducing FBXO44 later to prevent prolonged mutagenic synthesis once bypass is complete.

In cancer datasets, FBXO44 expression is associated with TP53 status, tumor mutational burden, therapy response, relapse-related features, and prognosis. These associations suggest that disruption of the p53-FBXO44-RAD18 axis may contribute to hypermutation and therapy adaptation, particularly in tumors with compromised p53 signaling.

In summary, FBXO44 provides a p53-associated layer of control over RAD18 and TLS termination. By limiting persistent PCNA monoubiquitination, FBXO44 helps restrain mutagenesis after replication stress. Our findings help us understanding the mechanims for tumor evolution under therapeutic pressure and how these could be exploited therapeutically.

## Supporting information

Suppelmentary Figures

## Funding

This work was supported by the Deutsche Krebshilfe to IA (#70117491), by the DFG to IA and AJG (TRR353 "Death Decision", project A05) and the Carl Zeiss Stiftung to IA (Endowed Professorship #15972218).

## Data availability

Data are available from the corresponding author upon reasonable request.

## Declarations

### Ethics approval and consent to participate

Not applicable.

### Human Ethics and Animal Ethics statements

Not applicable.

### Consent for publication

Not applicable.

## Availability of data and materials

Data and materials are available from the corresponding author upon reasonable request.

## Competing interests

The authors declare that they have no competing interests.

## Materials and Methods

### Cell culture, siRNA transfection, and treatments

The human cell lines A549, HEK293T, U2OS, HCT-116, and MDA-MB-231 were cultured in Dulbecco’s modified Eagle’s medium (DMEM; high glucose, GlutaMAX) supplemented with 10% fetal bovine serum (FBS) and penicillin-streptomycin. Human lung epithelial HBEC-3KT cells were cultured in RPMI supplemented with 10% FBS and penicillin-streptomycin. All cell lines were maintained at 37°C with 5% CO2 and were routinely tested for mycoplasma.

For silencing experiments, cells were transfected with 20 nM siCTR (Negative Control #1, Thermo Fisher Scientific, #4390844), p53 siRNA (Thermo Fisher Scientific, #4390825, ID s605), or FBXO44- targeting siRNAs (Thermo Fisher Scientific, #4392420; IDs s41168, s41169, s41170, and s225117) using Lipofectamine RNAiMAX (Invitrogen), according to the manufacturer’s protocol. Cells were collected 72 h after transfection. For p53 induction, transfected cells were treated with Nutlin-3a (10 µM) for the indicated times. For replication-stress experiments, A549 cells were treated 72 h after FBXO44 knockdown with 2 mM hydroxyurea (HU) for the indicated times and then allowed to recover after HU withdrawal.

### Chemicals

The MDM2 inhibitor Nutlin-3a was purchased from Selleckchem (S8059), resuspended in DMSO, and used at the indicated concentrations. Hydroxyurea (HU; H8627), cisplatin (232120), and 6- thioguanine (6-TG; A4882) were purchased from Sigma-Aldrich. HU and cisplatin were resuspended in cell-biology-grade H2O and diluted in medium as indicated. 6-TG was resuspended in 1 M NaOH and diluted in medium as described.

### Plasmids and transfection

The FBXO44 open reading frame (ORF; transcript variant 1) was subcloned from pCMV6-FBXO44 (Origene, RC216126) into pcDNA3.1 and tagged at the N terminus. The FBXO44 F-box construct was cloned from full-length FBXO44 into pcDNA3.1 and N-terminally tagged. The Myc-hRAD18 expression plasmid (Addgene #68827) was a gift from the Tateishi laboratory^27^. TurboID-V5-FBXO44 and BirA(WT)-V5-FBXO44 plasmids were generated using Gateway cloning.

HEK293T cells (3 × 10^6^) were seeded in 10-cm dishes 24 h before plasmid transfection. For each dish, 5 µg plasmid DNA and 30 µl PEI MAX (Polysciences, cat. no. 24765) were diluted in 500 µl Opti-MEM, vortexed, and incubated for 30 min at room temperature. The transfection mixture was then added dropwise. PEI stock (1 mg/ml) was prepared according to the manufacturer’s instructions and filter sterilized. Twenty-four hours after transfection, cells were washed twice with PBS and harvested. Cell pellets were lysed directly for downstream experiments.

### RNA extraction, reverse transcription, and qPCR

RNA was extracted using the RNeasy Mini Kit (Qiagen) according to the manufacturer’s instructions. A total of 1 µg RNA was reverse transcribed using the SensiFAST cDNA Synthesis Kit (Biocat, BIO- 65054). Real-time PCR (qPCR) was performed using PerfeCTa SYBR Green FastMix (VWR, 733- 1390). The following primers were used: FBXO44 forward, 5’-TGAATGGAGGCGATGAGTGG-3’; FBXO44 reverse, 5’-GCTCCTCCCAATACCCTTCG-3’; TP53 forward, 5’- CCCAAGCAATGGATGATTTGA-3’; TP53 reverse, 5’-GGCATTCTGGGAGCTTTCATCT-3’; CDKN1A forward, 5’-CCTGTCACTGTCTTGTACCCT-3’; CDKN1A reverse, 5’- GCGTTTGGAGTGGTAGAAATCT-3’; ACTB forward, 5’-AAAGACCTGTACGCCAACA-3’; ACTB reverse, 5’-CGGAGTACTTGCGCTCAG-3’. RT-qPCR results were expressed as relative gene expression using the 2^-ΔΔCt method after normalization to beta-actin (ACTB).

### Protein extraction and immunoblot analysis

Cells were lysed in RIPA buffer supplemented with protease and phosphatase inhibitors (Sigma-Aldrich). Proteins (5-20 µg) were heat denatured at 95°C for 10 min, resolved by SDS-PAGE, and transferred onto Amersham Hybond P 0.2 PVDF membranes (Sigma-Aldrich). Membranes were blocked for 1 h at room temperature in 5% non-fat dry milk in PBS containing 0.1% Tween-20. The following primary antibodies were incubated overnight at 4°C: rabbit anti-FBXO44 polyclonal antibody (Sigma-Aldrich, HPA003363), mouse anti-beta-actin monoclonal antibody (Sigma-Aldrich, A1978- 200UL), mouse anti-p53 monoclonal antibody (DO-1, Santa Cruz, sc-126), mouse monoclonal anti-GAPDH clone GAPDH-71.1 (Sigma-Aldrich, G8795-200UL), mouse anti-FLAG tag (M2 clone, Sigma-Aldrich, B3111-1MG), rabbit anti-Myc tag antibody (Cell Signaling Technology, #2278S), and mouse anti-Myc tag antibody [9E10] (Abcam, ab32). Membranes were incubated with appropriate HRP-conjugated anti-mouse or anti-rabbit secondary antibodies (Bio-Rad) for 1 h at room temperature. Chemiluminescence signals were detected using ECL Chemiluminescence Kit (Thermo Fisher Scientific, #32106X4) or SuperSignal West Dura Extended Duration Substrate (Thermo Fisher Scientific, #34075).

### Co-immunoprecipitation (co-IP)

For co-IP experiments, cells were washed twice with cold 1x PBS and lysed for 30 min at 4°C with rotation in lysis buffer containing 50 mM Tris-HCl (pH 7.5), 150 mM NaCl, 1 mM EDTA, 5 mM MgCl2, and 0.1% NP-40, supplemented with protease (Roche, #11836170001) and phosphatase inhibitor cocktails (Roche, #4906845001). Lysates were centrifuged for 15 min at 4°C and precleared for 1 h with Sepharose Protein G (Cytiva, GE17-0618-01). For semi-endogenous or exogenous co-IP, 0.5-1 mg protein was used per pull-down; for endogenous co-IP, 1.5 mg protein was used. After addition of the indicated antibodies (0.5-1 µg for semi-endogenous/exogenous co-IP; 1.5 µg FBXO44 antibody or IgG control for endogenous co-IP), pull-downs were incubated overnight at 4°C with gentle rotation. Immune complexes were isolated the following day by incubation with Protein G magnetic beads (Invitrogen, #10003D) for 2 h at 4°C. Beads were washed three times with lysis buffer, eluted in 1x Bolt LDS sample buffer (Thermo Fisher Scientific) supplemented with Bolt Reducing Agent, boiled at 95°C for 10 min, and resolved on 4-12% Bolt Bis-Tris Plus Mini Protein Gels (Thermo Fisher Scientific, NW00120BOX).

### Enrichment of diGly peptides

MDA-MB-231 cells transfected with siCTR or siFBXO44 were lysed in urea lysis buffer (20 mM HEPES, pH 8.0, 9 M urea) supplemented with phosphatase inhibitor cocktail (Cell Signaling Technology, #5870). After tryptic digestion and C18 reversed-phase extraction, ubiquitinated peptides were enriched using protein A agarose beads conjugated to the PTMScan Ubiquitin K-GG Remnant Motif Antibody (Cell Signaling Technology, #59322). After resin washing and elution, peptides were analyzed by LC-MS/MS. MS/MS spectra were searched with Comet^45^ through the Harvard Core platform against the latest UniProt Homo sapiens database, using mass accuracy tolerances of +/-50 ppm for precursor ions and 0.02 Da for product ions. Results were filtered using +/-5 ppm precursor- ion mass accuracy and the presence of the intended motif.

### Recombinant protein production of GST-UBA^Ubq^

The plasmid encoding the UBA domain of Ubiquilin-1 (Q9UMX0, isoform 1, amino acids 536-589) fused to an N-terminal GST tag and C-terminal hexahistidine tag in pGEX6P1 was provided by Prof. V. D’Angiolella. The plasmid was transformed into E. coli BL21(DE3) cells (Thermo Fisher Scientific, #C600003). Cells were cultured in TB medium at 37°C until OD600 reached 2. Recombinant UBAUbq expression was induced by adding 0.4 mM isopropyl beta-D-1-thiogalactopyranoside, followed by 18 h shaking at 20°C. Cells were harvested by centrifugation and lysed by sonication in binding buffer (50 mM sodium phosphate, pH 7.5, 300 mM NaCl, 10% glycerol, 10 mM imidazole, 0.5 mM TCEP). UBAUbq was captured on nickel Sepharose resin (Thermo Fisher Scientific, #88221), washed with binding buffer containing 30 mM imidazole, and eluted four times with elution buffer (50 mM sodium phosphate, pH 7.5, 300 mM NaCl, 10% glycerol, 500 mM imidazole, 0.5 mM TCEP). UBAUbq purity was assessed by Coomassie staining.

### MS sample preparation for TurboID-FBXO44 pull-down

For TurboID-MS experiments, procedures were performed as previously described^46^. Briefly, TurboID- FBXO44 Flp-In T-REx HEK293 cells were grown in 15-cm dishes to 80% confluency. Doxycycline (1.3 µg/ml) was added for 24 h to induce TurboID-FBXO44 expression, followed by incubation with 50 µM biotin for 3 h to label proximal proteins. Cells were harvested by scraping, washed three times with cold 1x PBS, and resuspended in 1 ml RIPA buffer (50 mM Tris-HCl, pH 8.0, 150 mM NaCl, 1% Triton X-100, 1 mM EDTA, and 0.1% SDS) supplemented with protease inhibitor cocktail (Sigma-Aldrich, P8340). Lysates were incubated on ice for 15 min and clarified by centrifugation. Cleared lysates were incubated with 50 µl pre-equilibrated Strep-Tactin Sepharose beads (IBA, 2-1208-002) for 1 h at 4°C with rotation.

The suspension was loaded onto Mini Bio-Spin columns (Bio-Rad, 732-6207) to collect the beads. Beads were washed twice with 1 ml RIPA buffer, three times with HNN buffer (50 mM HEPES, pH 7.5, 150 mM NaCl, and 50 mM NaF), and twice with 100 mM NH4HCO3.

### Beads were transferred to 2-ml Eppendorf tubes in 400 µl 100 mM NH4HCO3

After centrifugation at 200 × g for 1 min, beads were resuspended in 100 µl 8 M urea in 100 mM NH4HCO3 and incubated for 20 min at 20°C. Cysteine bonds were reduced with 5 mM TCEP for 30 min at 37°C and alkylated with 10 mM iodoacetamide for 30 min at room temperature in the dark. Beads were digested with trypsin/Lys-C Mix (Promega, V5071) at a 25:1 protein:protease ratio (w/w) for 4 h at 37°C with orbital shaking. Urea was diluted to 1 M by adding 100 mM NH4HCO3, and digestion continued overnight at 37°C. Samples were desalted on C18 spin columns (Thermo Fisher Scientific, 89870), washed according to the manufacturer’s instructions, and eluted with 0.1% trifluoroacetic acid (TFA) and 65% acetonitrile.

Peptides were dried in a SpeedVac vacuum concentrator and resuspended in 0.1% TFA and 2% acetonitrile in MS-grade water for MS analysis.

### Immunofluorescence and confocal microscopy

PCNA monoubiquitination (PCNA-mUb) and CPDs were detected as previously described^47^ with minor modifications. Cells were seeded on glass slides, irradiated with 20 J/m2 UV-C, and fixed at the indicated times with 90% methanol in 1x PBS for 15 min at room temperature. Fixed cells were blocked with 2% bovine serum albumin for 2 h at room temperature. The following primary antibodies were incubated overnight at 4°C: ubiquitinated PCNA (Lys164; Cell Signaling Technology, #13439) and CPDs (Cosmo Bio, #CAC-NM-DND-001). Slides were washed three times with 1x PBS and incubated with the appropriate Alexa Fluor secondary antibody (Thermo Fisher Scientific) and DAPI. After three washes with 1x PBS, slides were mounted with ProLong Gold Antifade mounting solution (Thermo Fisher Scientific, P36934) and acquired in Z-stack mode on a Zeiss LSM 700 confocal microscope using ZEN software. Images were processed in Fiji, and overlap was measured as the percentage of area covered by the two channels.

### Ubiquitin Binding Entities (UBE) pull-down assay

The GST-UBA ubiquitin-associated domain of UBQLN1 was conjugated to glutathione Sepharose beads (Cytiva, GE17-0756-01) in ubiquitin binding entities (UBE) lysis buffer [19 mM NaH2PO4, 81 mM Na2HPO4 (pH 7.4), 1% NP-40, 2 mM EDTA, protease and phosphatase inhibitor cocktail, 50 mM N-ethylmaleimide (NEM; Sigma-Aldrich, E3876), 5 mM 1,10-phenanthroline (Sigma-Aldrich, 131377), and 50 µM PR-619 (ApexBio, A821)] for at least 4 h at 4°C with rotation. For each pull-down, 100 µg recombinant GST-UBA was conjugated to 20 µl washed glutathione Sepharose beads. Cells were left untreated or treated with MG132 (10 µM, 4 h) or HU (2 mM, 1 h) before lysis. Cells were lysed in freshly prepared UBE lysis buffer for 30 min at 4°C with rotation and centrifuged at 14,000 rpm for 15 min at 4°C. Protein concentration was measured by Bradford assay (Bio-Rad, 5000006), and equal amounts of total protein were used for each pull-down (1-3 mg). GST-UBA-conjugated beads were added to lysates and incubated overnight at 4°C with rotation. Beads were collected by centrifugation, washed five times with UBE lysis buffer, mixed with 1x Bolt LDS sample buffer, and boiled at 95°C for 10 min with shaking. Supernatants were used for immunoblotting. Total ubiquitin was probed as a loading control for UBE pull-down experiments.

### E3-substrate tagging by ubiquitin biotinylation (E-STUB)

E-STUB assays were performed as recently described^33^ with minor modifications. HEK293T cells were grown in DMEM supplemented with 10% Tet-free FBS for 3-4 days before the experiment. Three million cells were seeded per 10-cm dish. The following day, cells were co-transfected with empty vector (EV) or BirA-V5-FBXO44 expression vector together with pRK5-HA-A3-Ubiquitin. After 24 h induction, 50 µM biotin was added for 3 h. Biotin-labeled ubiquitinated species were pulled down with Streptavidin Sepharose beads (Cytiva, 17-5113-01) for 2 h at 4°C. After five washes with E-STUB RIPA buffer, beads were heated in Bolt LDS sample buffer containing 2 mM biotin at 70°C for 15 min with shaking. Eluted proteins were resolved on Bolt Bis-Tris Plus Mini Protein Gels (Thermo Fisher Scientific).

### Cycloheximide (CHX) chase

Cells (1 × 10^6^ per condition) were seeded in 6-cm dishes. The following day, cells were treated with cycloheximide (CHX; 100 µg/ml) for the times indicated in the figures to block protein synthesis and assess protein stability. Cells were washed twice with cold 1x PBS and lysed directly in RIPA buffer. Protein half-life was estimated by immunoblotting.

### HPRT mutagenesis assay

A549 FBXO44 wild-type (FBXO44+/+) and knockout (FBXO44-/-) clones, as well as U2OS cells, were maintained in DMEM supplemented with HAT (0.1 mM sodium hypoxanthine, 0.4 µM aminopterin, 0.016 mM thymidine; Gibco, #21060017) for 2 weeks. Cells were then released into DMEM supplemented with HT (0.1 mM sodium hypoxanthine, 0.016 mM thymidine; Gibco, #11067030) for 7 days. Cells were either irradiated with 20 J/m2 UV-C (A549), treated with cisplatin (U2OS), or left untreated. After 7-14 days of subculture, cells were seeded at 2 × 10^5^ cells (A549) or 5 × 10^5^ cells (U2OS) per 10-cm dish (seven plates total) in DMEM containing 5 µg/ml 6-TG (Sigma). After 4 weeks, 6-TG-resistant colonies were fixed with paraformaldehyde, stained with crystal violet, and counted. Plating efficiency was calculated by seeding 1 × 10^3^ cells in DMEM without 6-TG in triplicate for 2 weeks and counting stained colonies. Mutation frequency was calculated by dividing colony number by the total number of seeded cells and normalizing to plating efficiency.

### Colony formation assay

For clonogenic assays under cisplatin treatment, A549 FBXO44+/+ and FBXO44-/- cells were seeded at 3 × 10^5^ cells/well in 6-well plates. The next day, cells were treated with 50 µM cisplatin for 24 h. JH-RE-06 was added for an additional 24 h after cisplatin treatment. After drug removal, wells were washed three times with 1x PBS and surviving cells were grown for 4 weeks. For plating efficiency, 1 × 10^3^ cells/well were seeded. After 1 week, control wells were fixed with PFA for 20 min at room temperature and stained with crystal violet for 20 min. Colonies were counted manually.

### Generation of FBXO44 knockout cell lines

The FBXO44-targeting guide RNA (sgRNA) was cloned into pSpCas9(BB)-2A-Puro (PX459) V2.0 (Addgene, #62988). To generate A549 and HEK293T knockout cells, the Cas9-sgFBXO44 vector was transiently transfected using Lipofectamine 3000, as described above. At 24 h after transfection, cells were trypsinized, centrifuged, and reseeded in the same dish in the presence of 1 µg/ml puromycin (Thermo Fisher Scientific, #J67236.8EQ). After 36 h, puromycin was removed and cells were grown as single colonies. Effective FBXO44 knockout was confirmed by immunoblot analysis. The FBXO44 sgRNA sequence was 5’-CACCGAGGATCTCTCTCGAGACCAG-3’.

### 3D modeling of the RAD18-FBXO44 interaction

To study FBXO44-RAD18 binding in silico, we first examined each protein separately. For FBXO44, the surface topography of the crystal structure (PDB 3WSO)^48^ was analyzed with UCSF Chimera^49^ to identify crevices and electrostatic clusters. For RAD18, available experimental structures were mapped to the sequence and complemented with predictions of secondary structure and intrinsically disordered regions using JPred4 (https://www.compbio.dundee.ac.uk/jpred)^50^, AIUPRED (https://aiupred.elte.hu)^51^, and the MESSA meta-server (http://prodata.swmed.edu/MESSA)^52^ (Fig. S1A). Helical elements within disordered regions were further validated using PEP-FOLD3 (https://bioserv.rpbs.univ-paris-diderot.fr/services/PEP-FOLD3)^53^, and amphipathicity was analyzed by wheel projection in NetWheels (http://lbqp.unb.br/NetWheels)^54^.

The RAD18-FBXO44 interaction was then modeled with AlphaFold3 (https://alphafoldserver.com)^30^ and HDOCK (http://hdock.phys.hust.edu.cn)^31^, and consensus across algorithms was evaluated. Three-dimensional models of protein complexes were calculated from sequence data using AlphaFold3 for FBXO44 with either full-length RAD18, the RAD18 N-terminal region (residues 14- 144), or a peptide fragment (residues 123-144) predicted by the initial full-length model to mediate binding. Because RAD18 dimerizes through its N-terminal RING-finger-containing region (PDB 2Y43)^55^, predictions involving full-length or N-terminal RAD18 were performed for both monomeric and dimeric assemblies. For each condition, the top five predicted models were retrieved, and consensus and per-residue confidence scores (predicted local distance difference test, pLDDT) were evaluated. HDOCK modeling used either the FBXO44 crystal structure (PDB 3WSO)^48^ or the corresponding AlphaFold3 model with the RAD18 segments described above. Binding interfaces were analyzed with PISA (https://www.ebi.ac.uk/pdbe/pisa)^56^ and PRODIGY (https://wenmr.science.uu.nl/prodigy)^57^.

### ChIP-seq analysis

Raw FASTQ files were downloaded from GSE262052^58^. Read quality was assessed with FastQC^59^ (v0.12.1). Paired-end adapters were trimmed with fastp^60^ (v1.0.1). Reads were aligned to the human hg38 reference genome using Bowtie2^61^ (v2.5.4). Duplicate alignments and alignments overlapping blacklisted regions were removed, and only reads with MAPQ >10 were retained. Coverage tracks were computed with deepTools^62^ bamCompare (v3.5.5) for each immunoprecipitation sample against the input control using --binSize 50, --normalizeUsing BPM, --scaleFactorsMethod None, -- smoothLength 50, and --operation log2. Peaks were called with MACS3^63^ (v3.0.3) using --keep-dup all.

### ATAC-seq analysis

Raw FASTQ files were downloaded from GSE262051^58^. Read quality was assessed with FastQC^59^ (v0.12.1). Paired-end adapters were trimmed with Trim Galore^64^ (v0.6.10) using --phred33 --paired -- 2colour 20. Reads were aligned to the human hg38 reference genome using Bowtie2^61^ (v2.5.4). Duplicate alignments were removed, and alignments with MAPQ >10 were retained. Nucleosome-free fragments (<=140 bp inserts) were selected in silico. ATAC-seq reads were strand-shifted (+4 nt forward, -5 nt reverse) to account for the Tn5 offset. Coverage tracks were computed with deepTools^62^ bamCoverage (v3.5.5) using --binSize 20 and --normalizeUsing BPM. Peaks were called with MACS3^63^ (v3.0.3) using --nolambda --keep-dup all. Peaks overlapping blacklisted regions were removed, and replicate peaks were merged.

### Bioinformatics

Clinical data for lung adenocarcinoma (LUAD) and other tumor types (PanCancer) were retrieved from cBioPortal (http://www.cbioportal.org)^65,66^. Patient cohorts were divided into two groups according to FBXO44 mRNA expression (low vs. high) or TP53 status (wild-type vs. mutant).

## Statistical analyses

Graphs and statistical analyses were generated with GraphPad Prism 9.0 (GraphPad Software Inc.). Unless otherwise indicated in figure legends, results are shown as mean +/- standard deviation. Statistical analyses for cBioPortal-derived data were performed using unpaired t-tests. Unless otherwise indicated, other statistical analyses were performed using ordinary one-way ANOVA. Kaplan-Meier analyses were evaluated with the Mantel-Cox test. All experiments were performed with at least three biological replicates unless otherwise indicated in the figure legends.

---

**Supplementary Figure 1.** RAD18 is a putative FBXO44 target. A) Immunoblot analysis showing FBXO44 depletion in RPE p53-/- cells used for clonogenic survival assays after AZD6738 treatment; one of three experiments is shown. B) Gene ontology analysis of enriched biological processes associated with FBXO44 loss. C) Normalized abundance of total ubiquitinated RAD18 and K201, K229, and K230 RAD18 peptides in FBXO44-proficient (siCTR) and FBXO44-deficient (siFBXO44) cells. N = 2 independent technical replicates. D) Co-IP showing interaction between endogenous RAD18 and Flag-FBXO44; one representative blot of three independent replicates is shown.

**Supplementary Figure 2.** FBXO44 interacts with the E3 ligase RAD18. Sequence annotation of RAD18 domains described by Notenboom et al.^67^ is highlighted on the protein sequence (salmon): RING finger, zinc finger, SAP DNA-binding domain, Rad6-binding domain (Rad6 BD), and Polη-binding domain (Polη BD). Secondary-structure elements from available experimental structures (PDB codes in square brackets) are shown in orange, with helices indicated as cylinders and beta strands as arrows. For regions lacking experimental structures, MESSA- predicted secondary structure is shown in grey. Regions of intrinsic disorder predicted by AIUPRED are indicated by a wavy line (disorder score >0.5). The RAD18 segment predicted here to bind FBXO44 is boxed in magenta.

**Supplementary Figure 3.** FBXO44 ubiquitinates RAD18 in vivo. A) Immunoblot analysis of RAD18 levels after ectopic FLAG-FBXO44 expression in the presence of the NAE inhibitor MLN4924. N = 3 independent biological replicates. B) UBE pull-down ubiquitination assay in the absence or presence of the proteasome inhibitor MG132. Upper panel shows whole-cell extract (WCE). N = 3 independent biological replicates. C) Cycloheximide chase assay in HEK293T FBXO44+/+ and FBXO44-/- cells after hydroxyurea (HU)-induced replication stress. N = 2 independent biological replicates. D) Cycloheximide chase assay in HEK293T cells transfected with pcDNA3.1 empty vector (EV) or pcDNA3.1-FLAG-FBXO44 after HU-induced replication stress. N = 2 independent biological replicates. For immunoblot analysis, one of two independent experiments is shown.

**Supplementary Figure 4.** FBXO44 loss associates with mutagenesis and chemoresistance. A) Micrograph of the HPRT assay in A549 FBXO44+/+ and FBXO44-/- cells after UV-C irradiation (20 J/m2). B) Clonogenic assay showing no difference in plating efficiency or proliferation capacity between A549 FBXO44+/+ and FBXO44-/- clones in relation to Fig. 3I, J. C) Immunoblot analysis showing endogenous FBXO44 silencing in U2OS cells; one of two independent experiments is shown. D) Violin plot showing disease-free status of LUAD patients stratified by FBXO44 mRNA expression. E) Violin plots showing that lower FBXO44 mRNA levels are associated with disease recurrence/progression; **p < 0.01. P values were calculated by unpaired t-test. F) Violin plots showing that lower FBXO44 mRNA levels are associated with the presence of tumor after treatment; *p < 0.05. P values were calculated by unpaired t-test. G) Kaplan-Meier analysis showing that lower FBXO44 expression is associated with worse disease-free survival. P values are indicated. Source for D-G: cBioPortal database.

**Supplementary Figure 5.** FBXO44 is a late p53 response gene. A) RT-qPCR (left) and immunoblot (right) for FBXO44, p53, and CDKN1A after p53 activation by Nutlin-3a in HBEC-3KT (p53+/+) cells. N = 3 independent biological replicates; *p < 0.05, **p < 0.01, ***p < 0.001, ****p < 0.0001. P values were calculated by one-way ANOVA with Sidak’s correction. B) RT-qPCR (left) and immunoblot (right) for FBXO44, p53, and CDKN1A after p53 activation by Nutlin- 3a in U2OS (p53+/+) cells. N = 3 independent biological replicates; *p < 0.05, **p < 0.01, ***p < 0.001, ****p < 0.0001. P values were calculated by one-way ANOVA with Sidak’s correction. C) RT-qPCR (left) and immunoblot (right) for FBXO44, p53, and CDKN1A after p53 activation by Nutlin-3a in HCT- 116 (p53+/+) cells. N = 3 independent biological replicates; *p < 0.05, **p < 0.01, ***p < 0.001, ****p < 0.0001. P values were calculated by one-way ANOVA with Sidak’s correction. D) RT-qPCR (left) and immunoblot (right) for FBXO44, p53, and CDKN1A after Nutlin-3a treatment in the p53-mutant (p53R280K) MDA-MB-231 cell line. N = 3 independent biological replicates; ns, not statistically significant; *p < 0.05, ****p < 0.0001. P values were calculated by one-way ANOVA with Sidak’s correction. For immunoblots, one of three independent experiments is shown.

**Supplementary Figure 6.** Replication stress induces p53-dependent upregulation of FBXO44. A) RT-qPCR for p53 and CDKN1A after UV-C treatment in HBEC-3KT (p53+/+) cells. N = 3 independent biological replicates; *p < 0.05, **p < 0.01, ***p < 0.001, ****p < 0.0001. P values were calculated by one-way ANOVA with Sidak’s correction. B) RT-qPCR for p53 and CDKN1A after UV-C treatment in A549 (p53+/+) cells. N = 3 independent biological replicates; *p < 0.05, **p < 0.01, ***p < 0.001, ****p < 0.0001. P values were calculated by one-way ANOVA with Sidak’s correction. C, D) RT-qPCR (C) and immunoblot (D) for FBXO44, p53, and CDKN1A after hydroxyurea-induced replication stress in A549 p53-proficient (p53+/+) and p53-deficient (p53-/-) cells. N = 3 independent biological replicates; *p < 0.05, **p < 0.01, ***p < 0.001, ****p < 0.0001. P values were calculated by one-way ANOVA with Sidak’s correction. For immunoblotting, one of three independent experiments is shown. E, F) Fbxo44 FPKM by RNA-seq (E) and validation by RT-qPCR (F), showing conserved regulation of Fbxo44 by p53 in the mouse pancreatic ductal adenocarcinoma (PDAC)-derived KpshC cell line. N = 3 independent biological replicates; *p < 0.05, ***p < 0.001. P values were calculated by unpaired t- test.

## Notes

### Competing Interest Statement

The authors have declared no competing interest.

