## Supplementary figures and images for "A p53-dependent FBXO44-RAD18 axis limits mutagenesis by terminating translesion DNA synthesis"

### Suppelmentary Figures

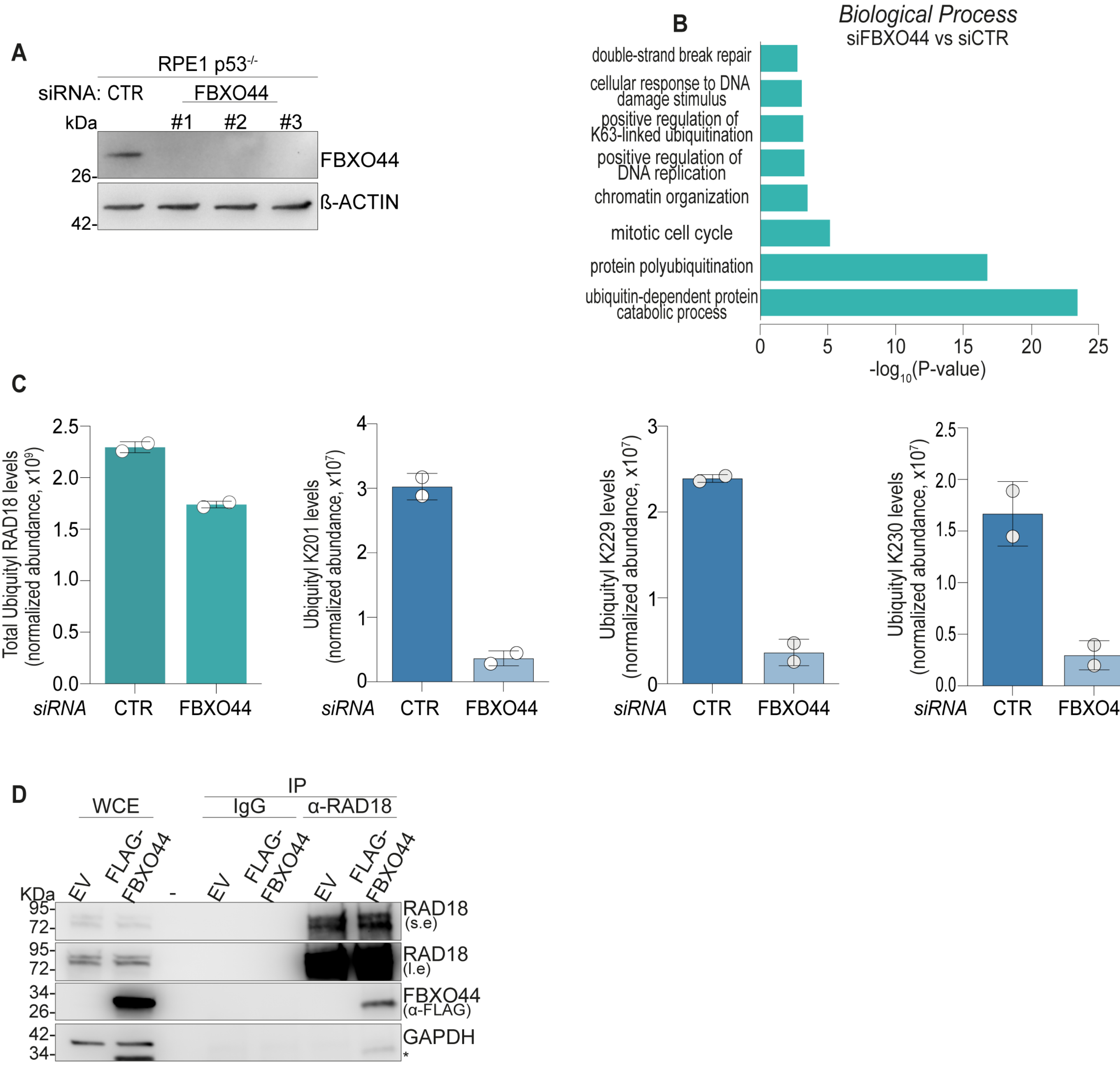


Suppl. Figure 1

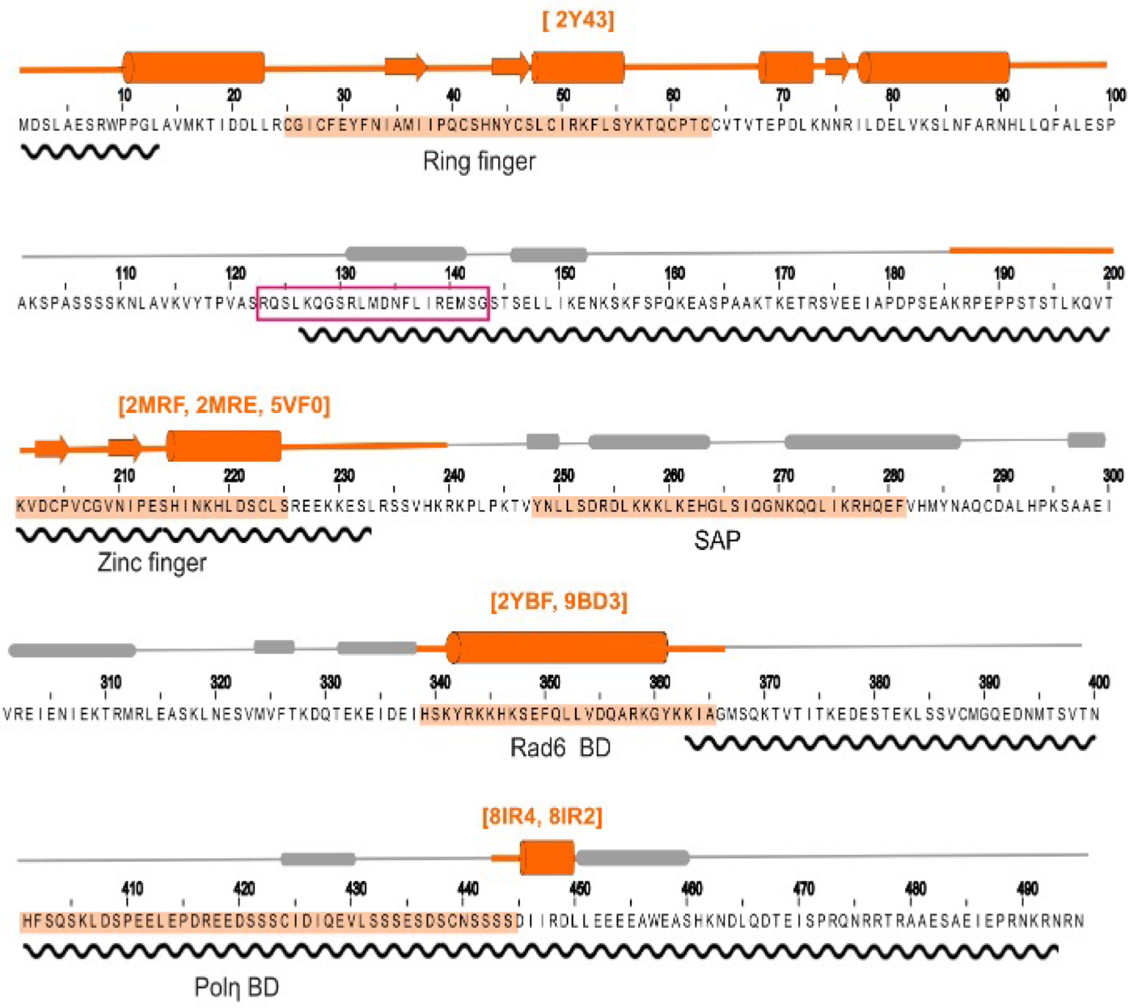


Suppl. Figure 2

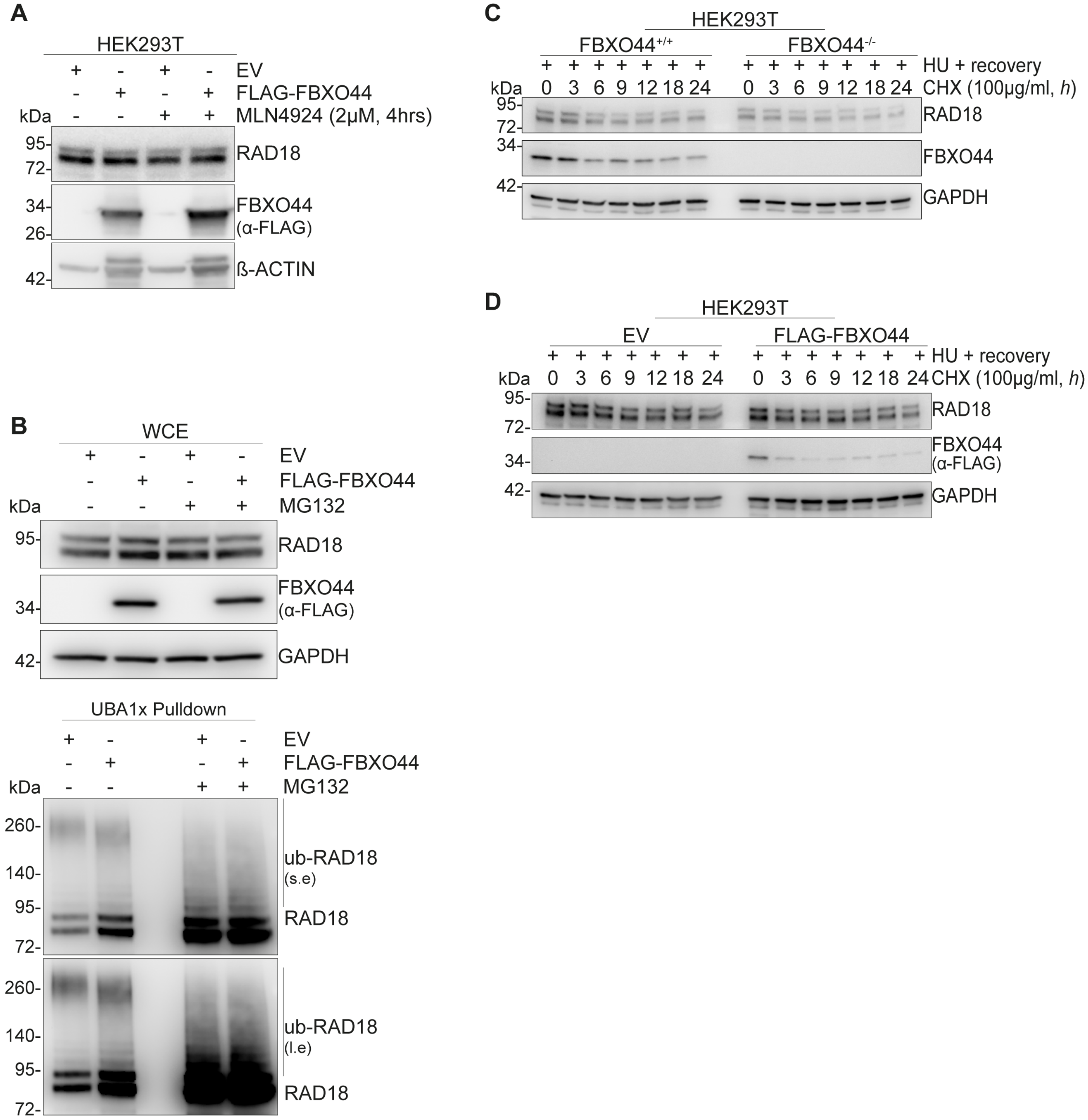


**Suppl. Figure 3**

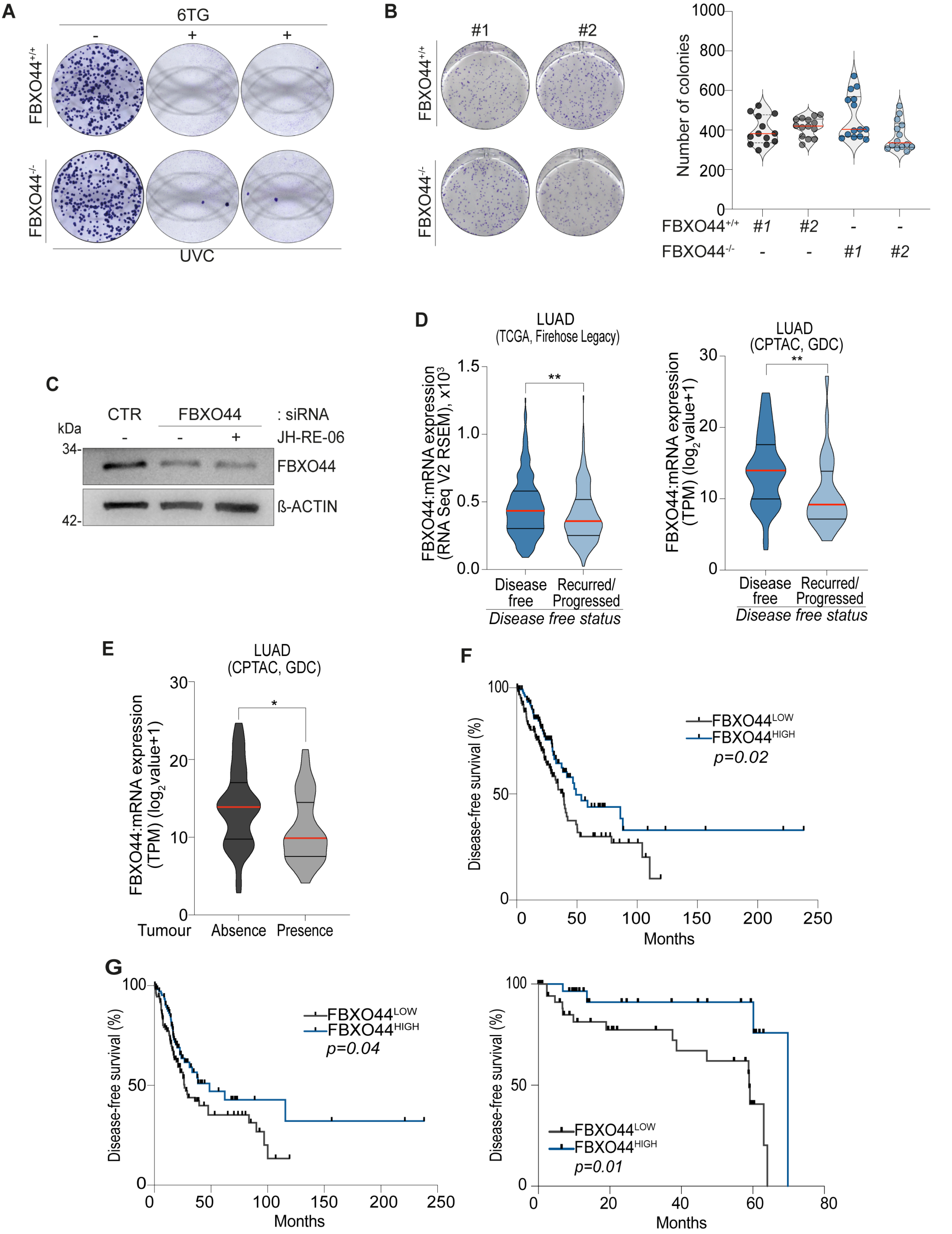


**Suppl. Figure 4**

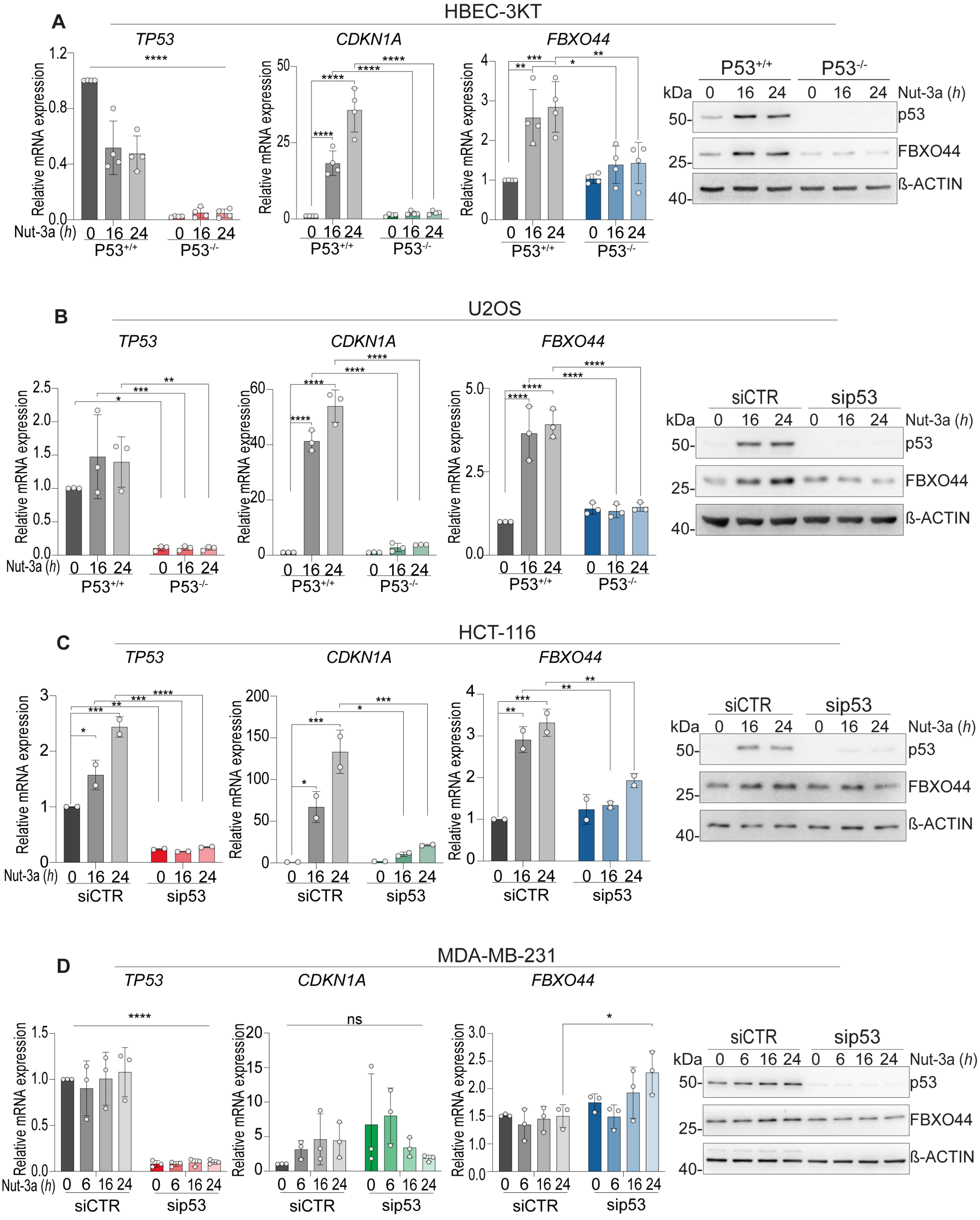


**Suppl. Figure 5**

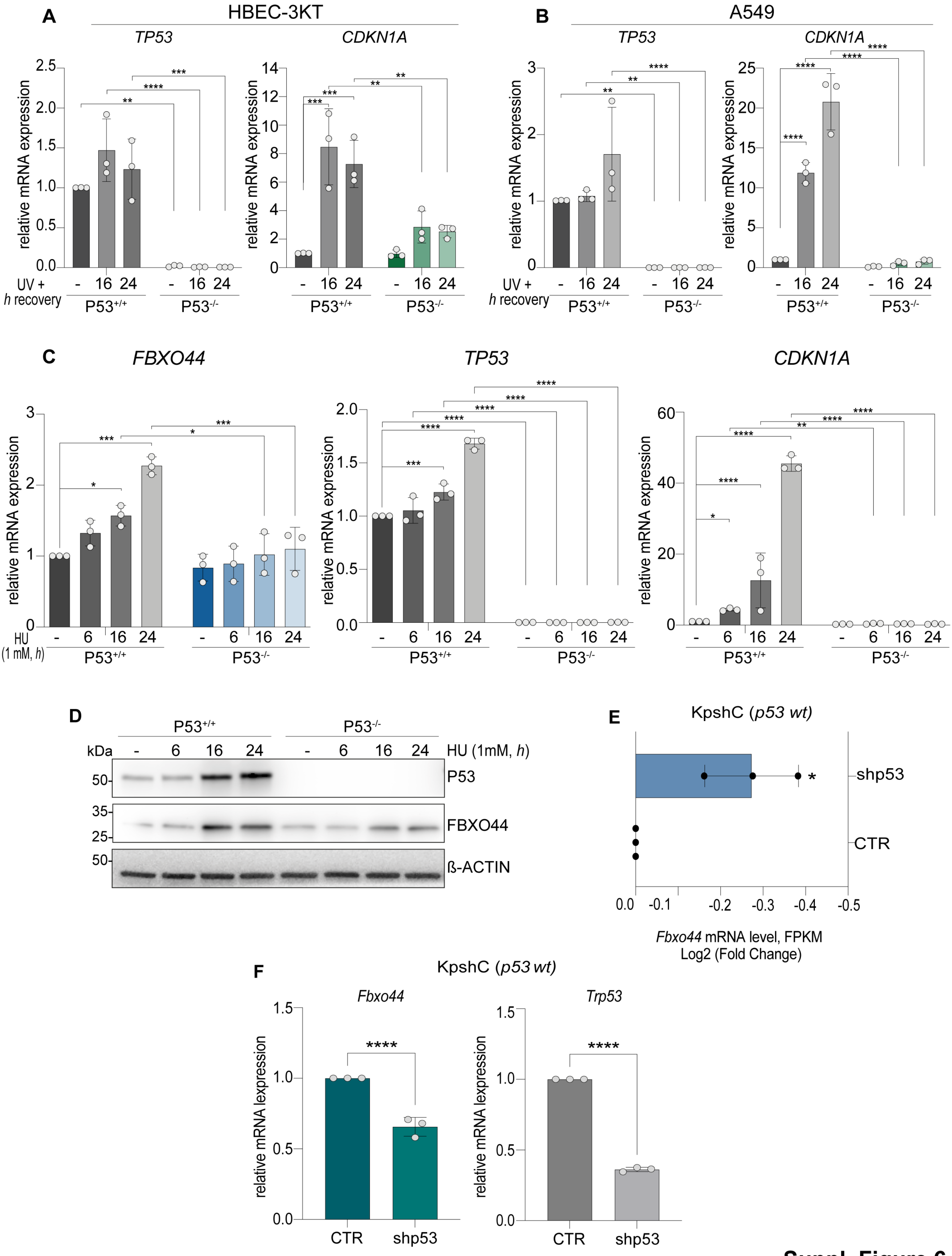


**Suppl. Figure 6**
